# CO-ADMINISTRATION OF RESVERATROL RESCUED LEAD-INDUCED TOXICITY IN *DROSOPHILA MELANOGASTER*

**DOI:** 10.1101/2023.03.09.532003

**Authors:** R. Abdulazeez, S. M. Highab, U.F. Onyawole, M.T. Jeje, H. Musa, D. M. Shehu, I. S. Ndams

**Affiliations:** Department of Zoology, Faculty of Life Sciences, Ahmadu Bello University, Zaria, Kaduna State, Nigeria; Department of Pharmacology and Therapeutic, Faculty of Basic Medical Sciences, College of Medicine and Health Sciences, Federal University Dutse, Jigawa State, Nigeria

**Keywords:** Fruit flies, Harwich strain, Lead poison, Resveratrol, Biological parameters, Oxidative stress

## Abstract

Lead toxicity poses a significant environmental concern linked to diverse health issues, including cognitive impairments, behavioral abnormalities, reproductive defects, and oxidative stress at the cellular level. This study explores the potential mitigating effects of resveratrol on lead-induced toxicity in *Drosophila melanogaster*. Adult *D. melanogaster* of the Harwich strain, aged three days, were orally exposed to lead (60 mg/L), Succimer (10 mg/kg), and varying doses of resveratrol (50, 100, and 150 mg/kg). The investigation encompassed the assessment of selected biological parameters, biochemical markers (ALP, AST, TB, CB, Na, Ca, Ur, Cr), oxidative stress indicators (MDA), and antioxidant enzymes (SOD and CAT). Resveratrol exhibited a dose-dependent enhancement of egg-laying, eclosion rate, filial generation output, locomotor activity, and life span in *D. melanogaster*, significantly to 150 mg/kg of diet. Most of the investigated biochemical parameters showed significant rescue in lead-exposed fruit flies when co-treated with resveratrol (p < 0.05). However, oxidative stress, as indicated by MDA levels, remained unaffected by resveratrol in this study. The findings suggest that resveratrol effectively protects against lead toxicity in *Drosophila melanogaster* and may hold therapeutic potential as an agent for managing lead poisoning in humans.

## Introduction

Heavy metal pollution, implicated in various diseases including kidney dysfunction (Tsai *et al*., 2017), nervous system disorders, and cardiovascular diseases (Obeng-Gyasi *et al*., 2018), poses a significant environmental and public health concern. Unlike organic matter, heavy metals such as lead (Pb) accumulate in organs like the kidney, liver, brain, and bone, leading to chronic poisoning and potentially fatal outcomes (Rehman *et al*., 2018; Cheng *et al*., 2019). The widespread dispersion of lead in the environment through industrial emissions and the use of lead-based products underscores the urgent need to address its impact on human health (CDC, 2023). Exposure to lead has been linked to reproductive toxicity, neurological damage, cardiotoxicity, oxidative stress, and reduced lifespan in both vertebrates and invertebrates (Flora *et al*., 2012; Rehman *et al*., 2018; Cheng *et al*., 2019).

Lead toxicity is particularly alarming for its association with cognitive deficits, Attention Deficit Hyperactivity Disorder (ADHD), altered sensory function, and delays in the onset of sexual maturity in children (Buchanan *et al*., 1999; Selevan *et al*., 2003; Braun *et al*., 2006; Jones and Miller, 2008; Nevin, 2009). In adults, lead poisoning contributes to cognitive impairment, cardiovascular and kidney dysfunction, and reproductive defects, with potential hereditary consequences for offspring (Winder, 1993; Perlstein *et al*., 2007; Kosnett *et al*., 2007; Kasperczyk *et al*., 2008; Weuve *et al*., 2009). Globally, lead poisoning accounts for 21.7 million disability-adjusted life years (DALYs) and deaths, with notable prevalence in countries like Nigeria and Senegal (W.H.O., 2023). The bioaccumulation of lead, affecting the hypothalamus– pituitary–gonadal axis, disrupts cellular functions and endocrine mechanisms, leading to inhibition of the mitochondrial electron transfer chain and the generation of reactive oxygen species (ROS) with genotoxic consequences (Gurer and Ercal, 2000; Helmut *et al*., 2010). Recognized as a serious public health issue, lead poisoning is conventionally treated with chelation therapy involving expensive drugs such as succimer (Vahabzadeh *et al*., 2021). In light of this, exploring readily available and cost-effective alternatives becomes imperative.

Studies indicate that certain polyphenol-producing plants exhibit significant antitoxic effects on lead toxicity (Gülçin, *et al.,* 2009; Gerszon, *et aal.,* 2014; Xia, *et al.,* 2017; Colica, *et al.,* 2018; Salehi, *et al.,* 2018; Gu, *et al.,* 2021; Hu, *et al.,* 2022; Rosiak, *et al.,* 2022). Resveratrol, a natural polyphenol found in grapes, wine, peanuts, and soy, has garnered attention for its antioxidant, anti-carcinogenic, anti-inflammatory, anti-aging, and anti-genotoxic properties (Burns *et al*., 2002; Baur *et al*., 2006; Yeh *et al*., 2013; Zhang *et al*., 2020).

*Drosophila melanogaster*, known as the vinegar fruit fly, emerges as a crucial invertebrate model in ecotoxicological research, serving as a reliable bioindicator of environmental pollution (Editorial team, 2023). The fruit fly shares similarities with humans in specific cell signaling pathways and protein-coding genes, such as metallothioneins protein (MTs), known for binding heavy metals (Al-momani and Massadeh, 2005). With its role as a predator, prey, pollinator, and decomposer, the fruit fly may serve as an entry point for lead into the terrestrial food web, making it an ideal model for studying the impacts of lead and resveratrol on various parameters (Skevington and Dang, 2002; Chifiriuc *et al*., 2016; Budiyanti *et al*., 2022). Moreover, the fruit fly’s affordability, ease of culturing, and short life cycle aligns with recommendations from the European Centre for the Validation of Alternative Methods (ECVAM) to promote the 3Rs (reduction, refinement, and replacement) in toxicity studies and testing (Benford *et al*., 2000).

Oxidative stress emerges as a crucial focal point in the exploration of lead toxicity (Obeng-Gyasi, 2018). Resveratrol, renowned for its capacity to neutralize reactive oxygen species, has been extensively documented in scientific literature (Leonard et al., 2003; Chandrashekara and Shakarad, 2011; Budiyanti et al., 2022). Notably, the compound’s widespread availability and affordability, even in resource-limited communities, position it as a potential cost-effective remedy for lead poisoning. Consequently, this investigation aims to assess the therapeutic potential of resveratrol in ameliorating the toxic effects induced by lead exposure in *Drosophila melanogaster*. The outcomes of this study may carry implications for comprehending the broader applicability of resveratrol in mitigating heavy metal toxicity and could contribute to the formulation of alternative and economical strategies for the treatment of lead poisoning.

## Materials and methods

### Chemicals

All chemicals were of analytical grade. *Trans*-resveratrol (60g) was obtained from Candlewood Stars Incorporated, Danbury, USA (Batch Number: MR 110218), Succimer (Sigma-Aldrich, U.S.A), while lead acetate (LA) (product No; 10142, BDH Laboratory Chemicals Limited Poole, England).

### Drosophila melanogaster stock and culture

The Harwich strain of *D. melanogaster* was obtained from Dr. A. Abolaji at the University of Ibadan and cultured in the *Drosophila* and Neurogenetics Lab within the Department of Zoology at Ahmadu Bello University, Zaria. The culture medium consisted of cornmeal with 1% w/v agar, 1% w/v Brewer’s yeast, 6% w/v corn flour, and 0.1% nipagin. The flies were maintained under consistent environmental conditions, including a temperature of 23 ± 2°C; 60% relative humidity, and a 12-hour dark/light cycle.

### Evaluating biological parameters

The egg-laying capability, filial generation output, and lifespan of flies subjected to resveratrol and lead acetate treatments were assessed following the protocols outlined by Chattopadhyay *et al*. (2015). The evaluation of the eclosion rate was conducted using the methodology described by Rand *et al*. (2014), and the negative geotaxis assay was implemented as per the procedure of Ali *et al*. (2011).

### Determination of biochemical and antioxidant parameters

To evaluate biochemical parameters, 50 flies of both genders were subjected to concentrations of lead acetate (60 mg/L), resveratrol (50, 100, and 150 mg/kg), and succimer (10 mg/kg) in 30 mg of diet over a 28-day period. Following the exposure period, the flies were anesthetized using CO_2_, weighed, and homogenized in 0.1 M phosphate buffer (pH 7.0) at a ratio of 1 mg to 10 mL. The homogenates were then centrifuged at 4000 g for 10 minutes at 4°C in a Thermo Scientific Sorval Micro 17 R refrigerated centrifuge. Subsequently, the supernatants were carefully separated from the pellets, transferred to labeled Eppendorf tubes, and stored at -20°C.

Analysis of liver function (fat body of the fruit flies) enzymes (AST, ALP, CB, and TB), serum electrolytes (Ca, Cl, Na, and K) and kidney function (Malpighian tubule) parameters (Ur and Cr) were conducted using an automated biochemistry analyzer (Selectra XL, Vital Scientific, Netherlands) and ELITech reagent kits from the Netherlands. Malondialdehyde activity was assayed following the method described by Niehaus and Samuelson, (1968), while catalase activity was determined according to the procedure outlined by Aebi, (1984). Superoxide dismutase activity was assessed using the method of Fridovich, (1989). All assays were performed in triplicate to ensure reliability and consistency of the results.

### Statistical analyses

The data were subjected to Levene’s homogeneity of variance test and Shapiro-Wilk test for normality before employing parametric statistics. One-way ANOVA was used to assess the significant differences among groups. Plots were made using the “ggplot2” package for R, and all statistical analyses were performed using R version 4.2.2 for macOS at a 5% significance level.

## Results

### Effects of lead toxicity and resveratrol on selected biological parameters of D. melanogaster

Lead exposure at 60 mg/L markedly decreased egg laying (fecundity). However, the detrimental effect of Pb was mitigated in a dose-dependent manner by resveratrol, with flies raised on 150 mg/kg of resveratrol exhibiting the highest egg-laying count (p < 0.05) compared to other treatments and the control group. Notably, the standard Pb poison treatment, succimer, did not restore the egg-laying ability of the lead-exposed fruit flies (see Figure 1). The eclosion rate (see Figure 2) was significantly diminished in flies exposed to lead. Resveratrol at a concentration of 50 mg/kg notably enhanced the emergence rate in multiple studied generations.

**Fig. 1.**
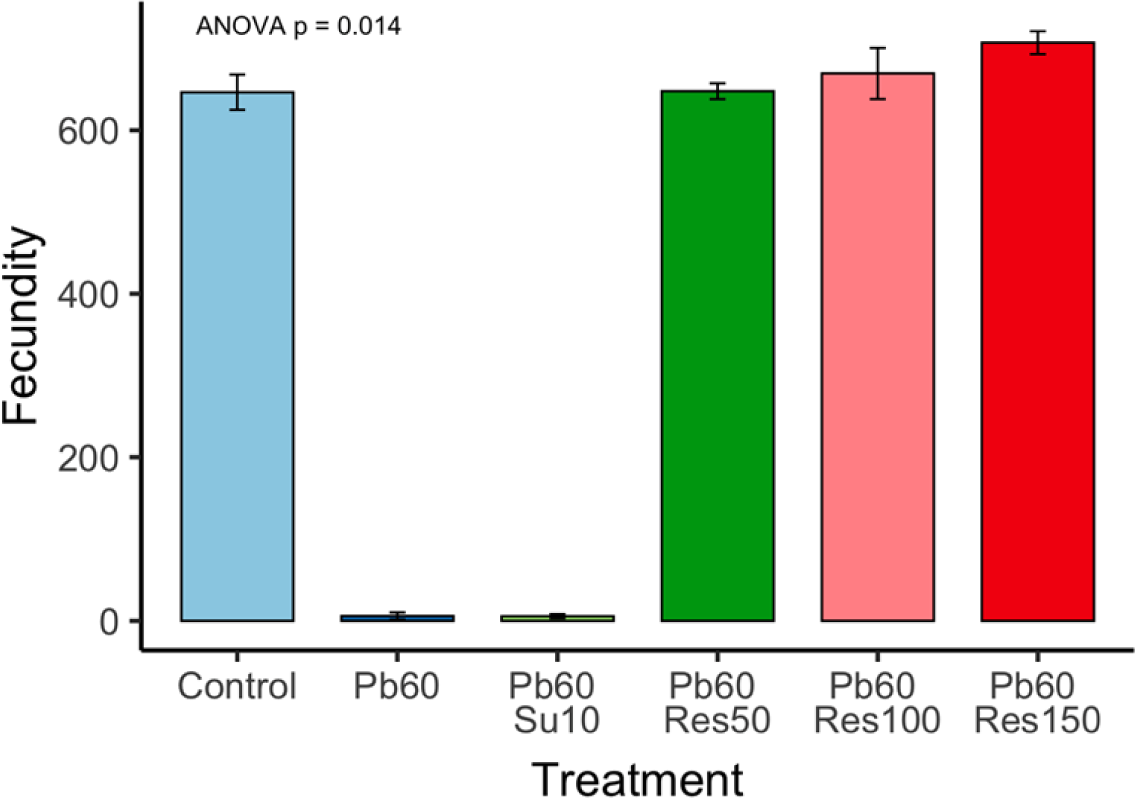
Impact of lead (Pb) and resveratrol on the egg-laying capacity of Harwich strain *D. melanogaster*. Data are expressed as Mean ± SEM per treatment group, with significance denoted as p < 0.05. Pb: lead concentration (mg/L), Su: succimer dosage (mg/kg), Res: resveratrol dosage (mg/kg)

**Fig. 2.**
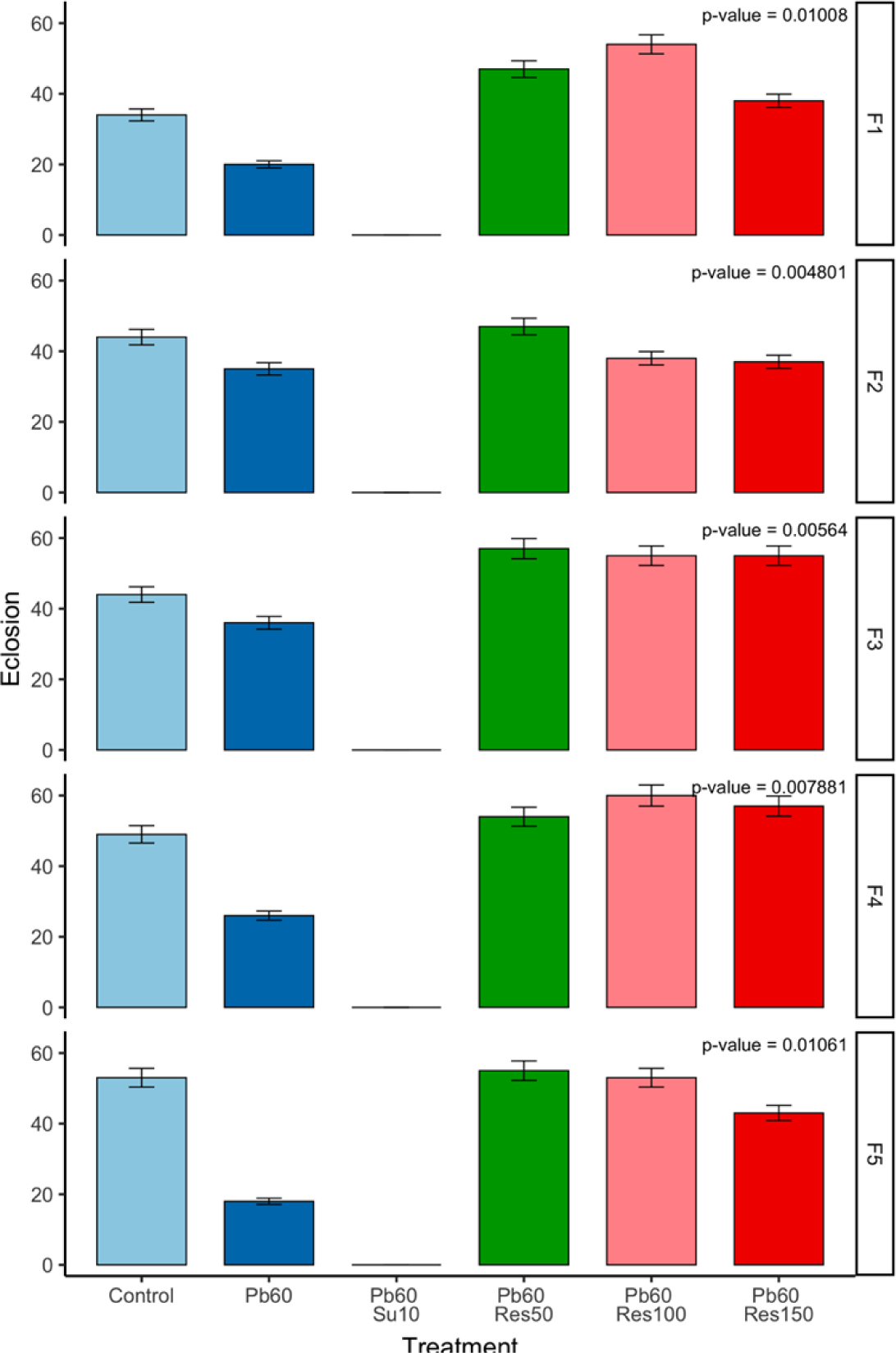
Implications of Pb and resveratrol on eclosure rate of Harwich strain *D. melanogaster.* Data are displayed as Mean ± SEM of 20 larvae/vial with 3 replicates per treatment group, and significance is indicated as p < 0.05. Pb: lead concentration (mg/L), Su: succimer dosage (mg/kg), Res: resveratrol dosage (mg/kg).

Analyzing the offspring production over five (5) generations, flies exclusively exposed to Pb exhibited a noteworthy transgenerational decline in filial generation output (see Figure 3), a reduction substantially alleviated by resveratrol at a concentration of 100 mg/kg in most generations. Furthermore, the locomotor activity for each filial generation (see Figure 4) revealed a marked decrease in the climbing ability of flies exposed to Pb. Notably, the treatment group with 150 mg/kg resveratrol significantly restored the climbing abilities of the flies. The lifespan of flies experienced a significant reduction upon exposure to Pb (34 days; p < 0.05), a decline that was markedly ameliorated to 53 days when co-exposed to Pb and 100 mg/kg resveratrol (see Figure 5; p < 0.05).

**Fig. 3.**
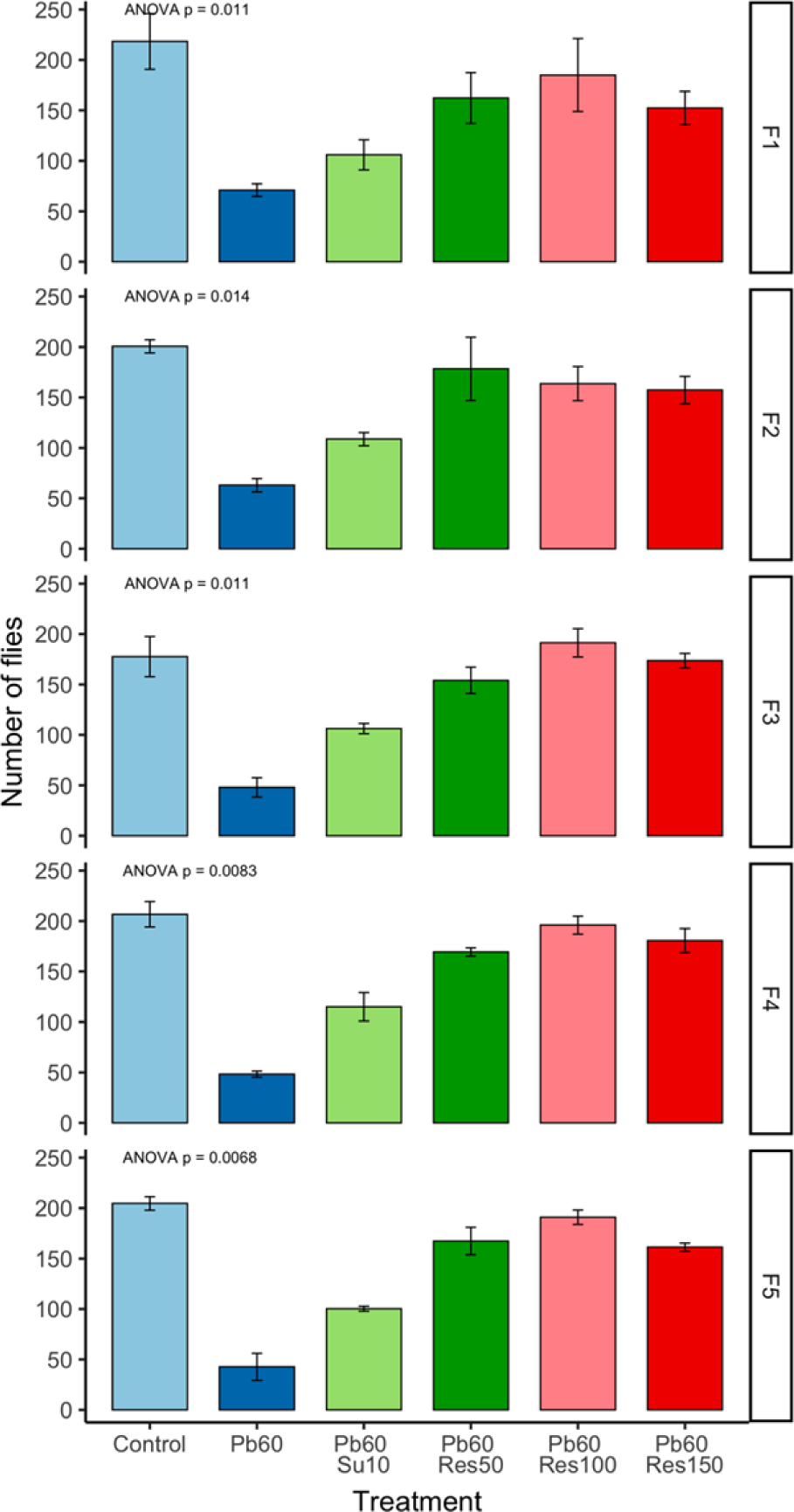
The impact of lead (Pb) and resveratrol on the filial generation output of Harwich strain *D. melanogaster* is illustrated. Data are expressed as Mean ± SEM of the number of emerged flies/vial, with 3 replicates per treatment group, and significance is denoted as p < 0.05. Pb: lead concentration (mg/L), Su: succimer dosage (mg/kg), Res: resveratrol dosage (mg/kg).

**Fig. 4.**
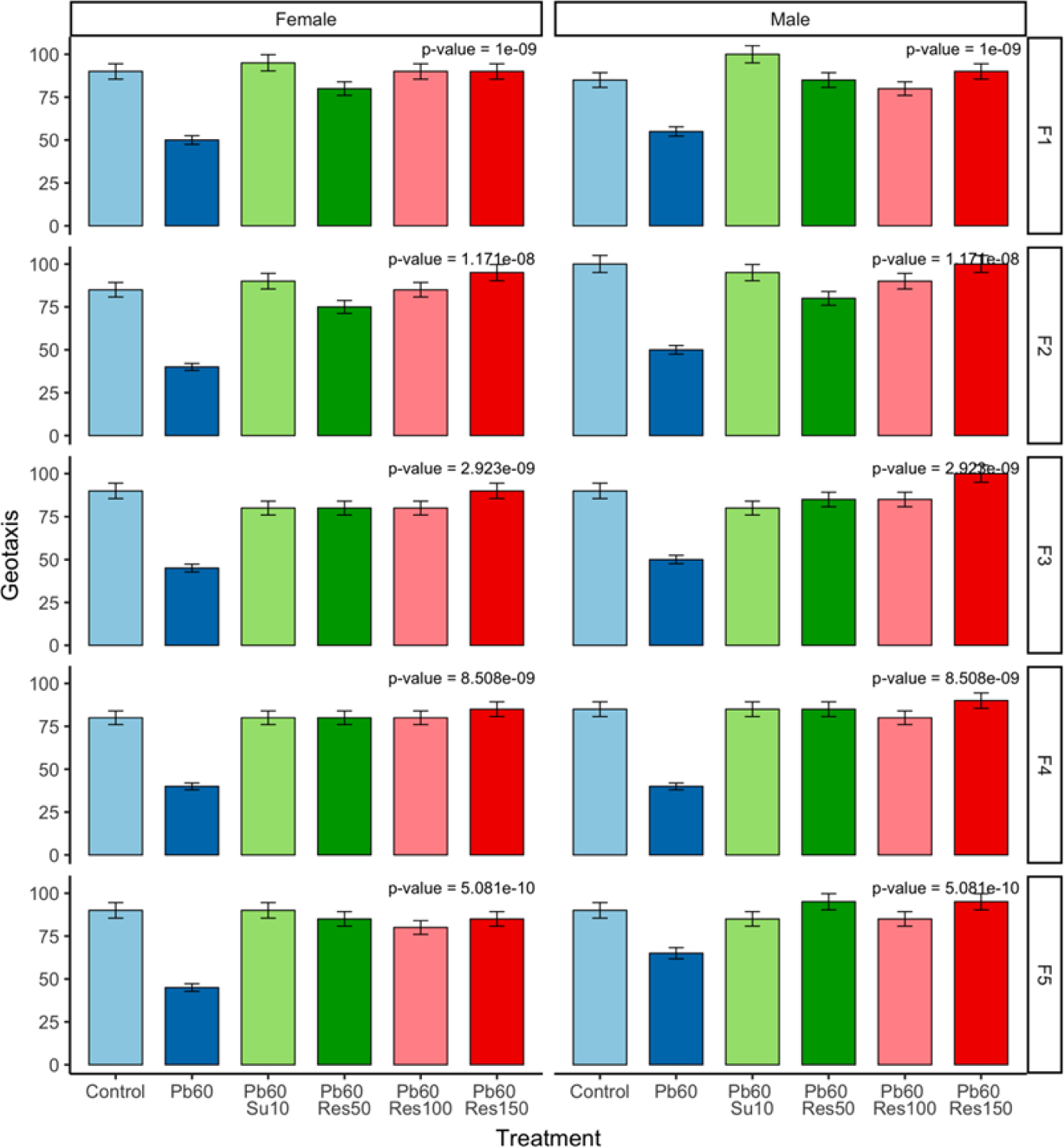
Effects of Pb and Res. on the climbing ability of male and female Harwich strain *D. melanogaster* is examined. Data are represented as the percentage of flies surpassing the 8 cm mark, presented as Mean ± SEM of 10 flies per 10 trials for each treatment group/generation, and significance is indicated as p < 0.05. Pb: lead concentration (mg/L), SU: succimer dosage (mg/kg), Res: resveratrol dosage (mg/kg).

**Fig. 5.**
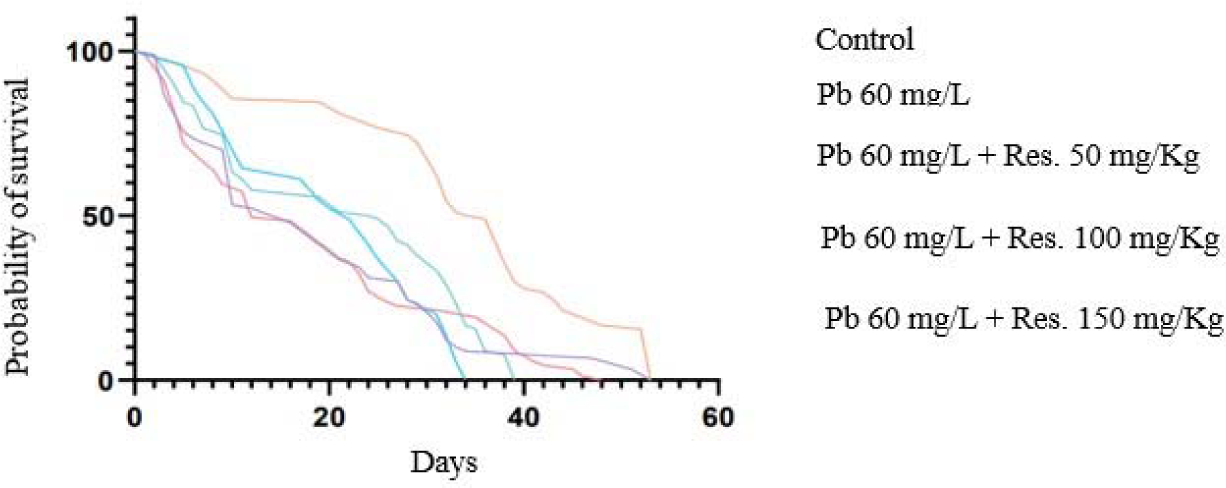
Influence of Pb and resveratrol on lifespan of *D. melanogaster* Harwich strain. Data are provided as the Mean of 50 flies/vial in triplicate for each treatment group, with significance denoted as p < 0.05. Pb: lead concentration (mg/L), Su: succimer dosage (mg/kg), Res: resveratrol dosage (mg/kg).

### Roles of lead and resveratrol on selected biochemical parameters of lead poisoned D. melanogaster

Prolonged exposure to lead (Pb), either independently or in conjunction with resveratrol, induced alterations in the examined biochemical parameters, enzymes, co-enzyme, electrolytes, oxidative stress markers, and antioxidants. The Pb-exposed group exhibited a noteworthy elevation in urea (Ur), total bilirubin (TB), and conjugated bilirubin (CB) compared to the control (p < 0.05). The level of Ur decreased in flies co-exposed to Pb at 60 mg/L + succimer (Su) at 10 mg/kg, while TB and CB decreased in flies co-exposed to Pb at 60 mg/L + resveratrol (Res) at 150 mg/kg. Conversely, the creatinine level was reduced in the presence of Pb alone and significantly increased in the Pb 60 mg/L + Res 50 mg/kg group (see Figure 6A-D).

**Fig. 6:**
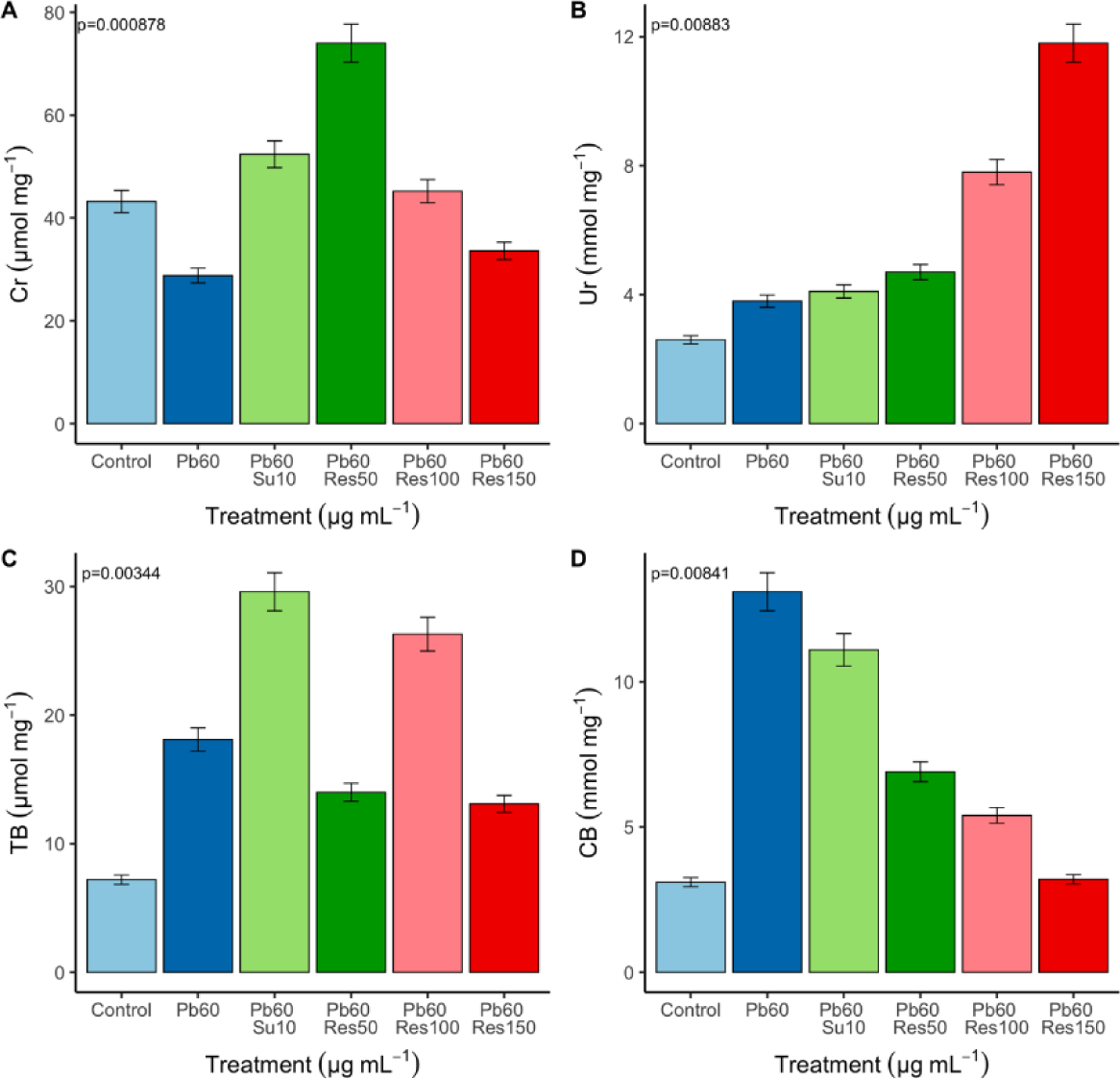
Effect of lead and resveratrol on the kidney and liver functions of lead-exposed *D. melanogaster.* Data are expressed as Mean ± SEM, with significance denoted as p < 0.05. Pb: lead concentration (mg/L), Su: succimer dosage (mg/kg), Res: resveratrol dosage (mg/kg)

In Figure (7A and C), a notable rise in the levels of the enzyme AST and co-enzyme Zn is evident in the Pb-treated group. Conversely, in the Pb 60 mg/L + Res 150 mg/kg group, a marked decrease in the levels of both enzymes is observed. The Pb-treated group displayed a significant reduction in the enzyme ALP concentration compared to the control group. In contrast, the Pb 60 mg/L + Res 150 mg/kg group exhibited a noteworthy increase in the ALP level (see Figure 6B; p < 0.05).

**Fig. 7:**
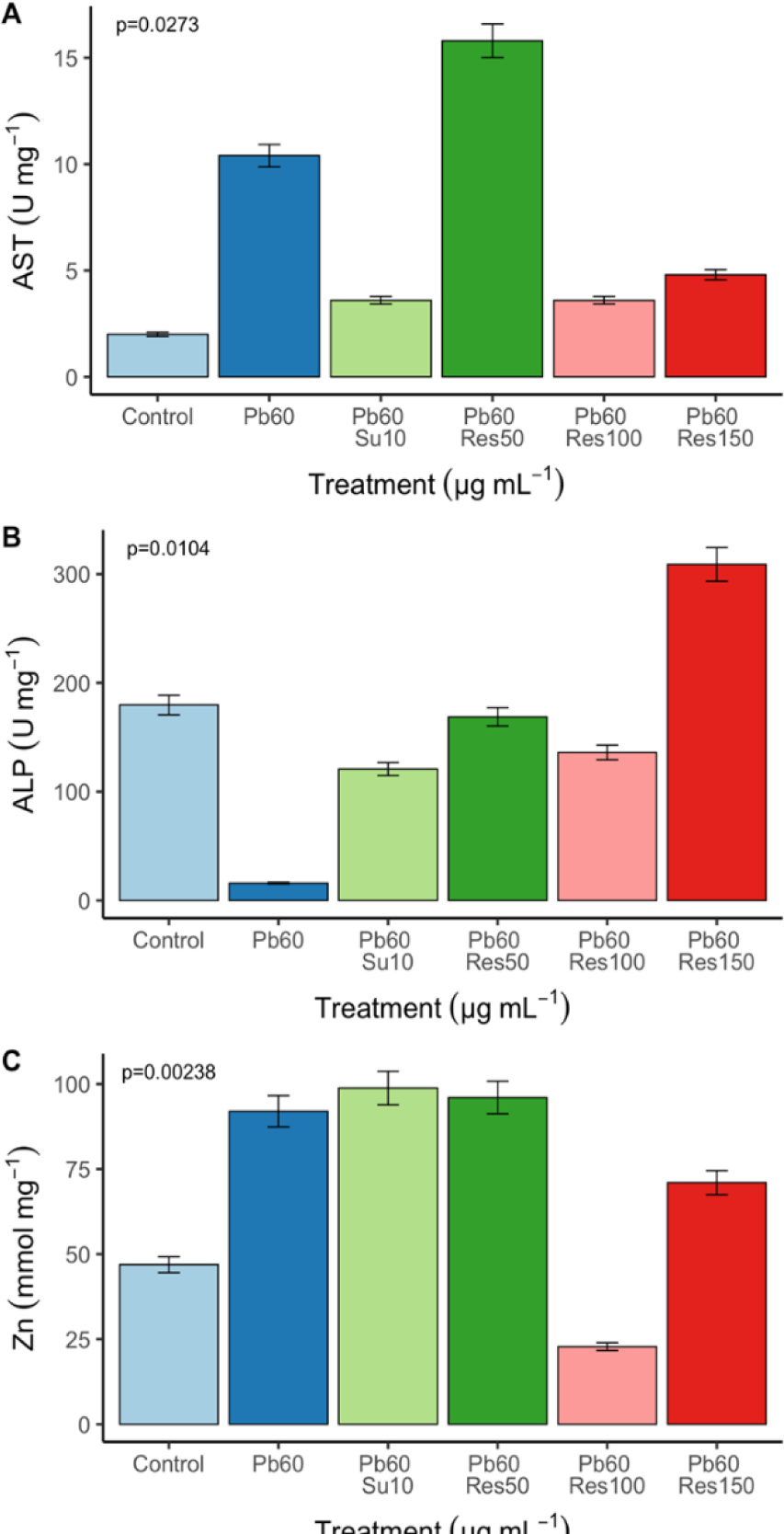
Effect of lead and resveratrol on serum biochemical parameters of lead-exposed *D. melanogaster.* The data is represented as Mean ± SEM, with significance indicated as p < 0.05. Pb: lead concentration (mg/L), Su: succimer dosage (mg/kg), Res: resveratrol dosage (mg/kg).

Alterations in the electrolytes Ca and K tended towards a decrease in the Pb-treated group, but statistical significance was noted in the Pb 60 mg/L + Res 150 mg/g group for both electrolytes (see Figure 8A and B). The Pb-treated group displayed an elevation in the Cl and Na levels, whereas reductions were observed in the Pb 60 mg/L + Res 150 mg/g group and Pb 60 mg/L + Res 100 mg/g group for Cl and Na, respectively (see Figure 8C and D).

**Fig. 8:**
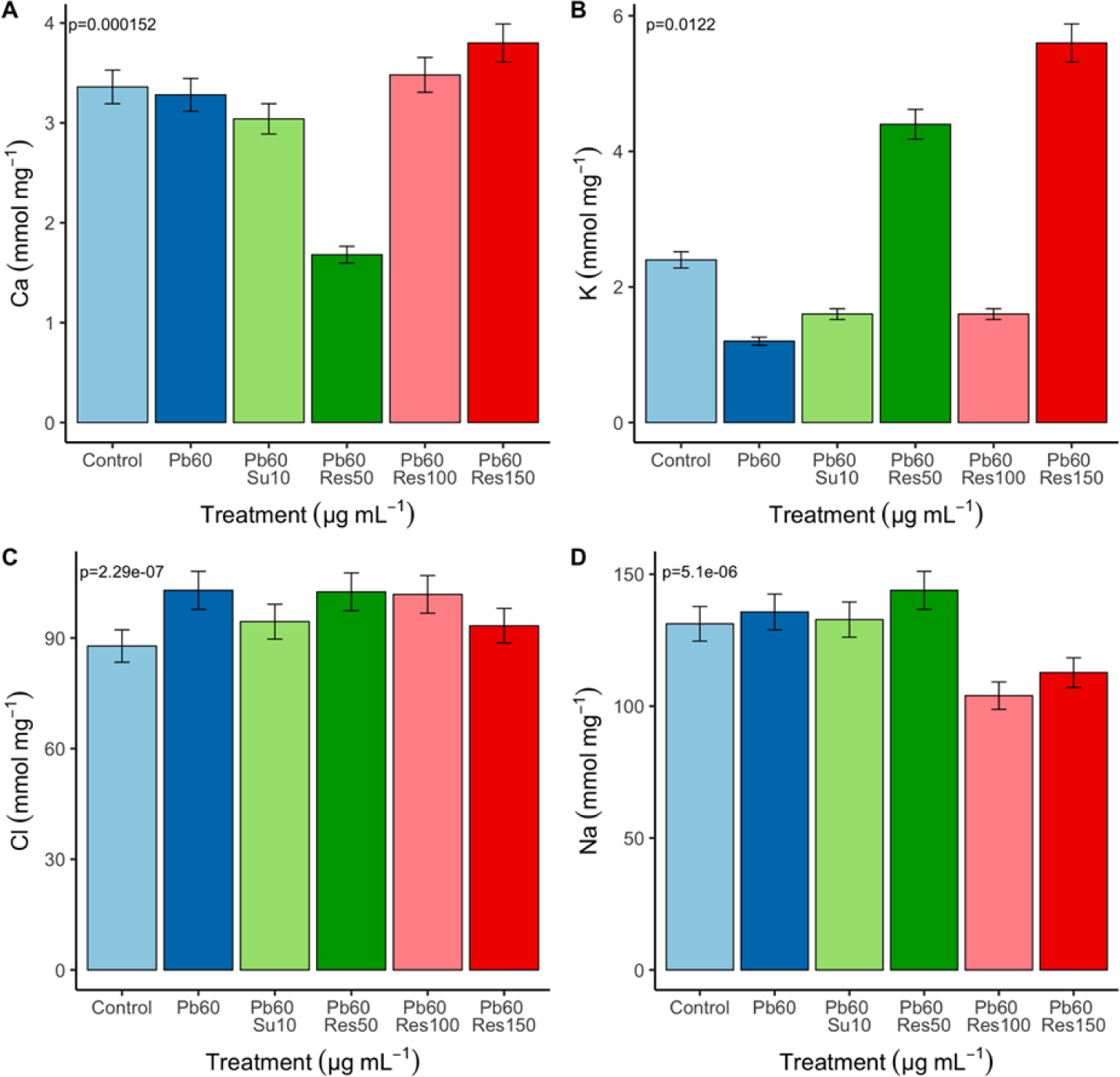
The impact of lead and resveratrol on the serum electrolytes of treatment groups is illustrated. Data are expressed as Mean ± SEM, with significance indicated as p < 0.05. Pb: lead concentration (mg/L), Su: succimer dosage (mg/kg), Res: resveratrol dosage (mg/kg).

All experimental groups exhibited an elevation in MDA and SOD levels in comparison to the control. However, a non-significant reduction (p > 0.05) in the oxidative stress marker and a noteworthy increase (p < 0.05) in the antioxidant were observed in the Pb 60 mg/L + Res 100 mg/g group (see Figure 9A and B). The activities of CAT in flies exposed to Pb and resveratrol were significantly different in all treated groups when compared with the normal control (p < 0.05). Additionally, the Pb 60 mg/L + Res 50 mg/g group displayed a significant increase in the CAT levels compared to other treated groups (see Figure 9C).

**Fig. 9:**
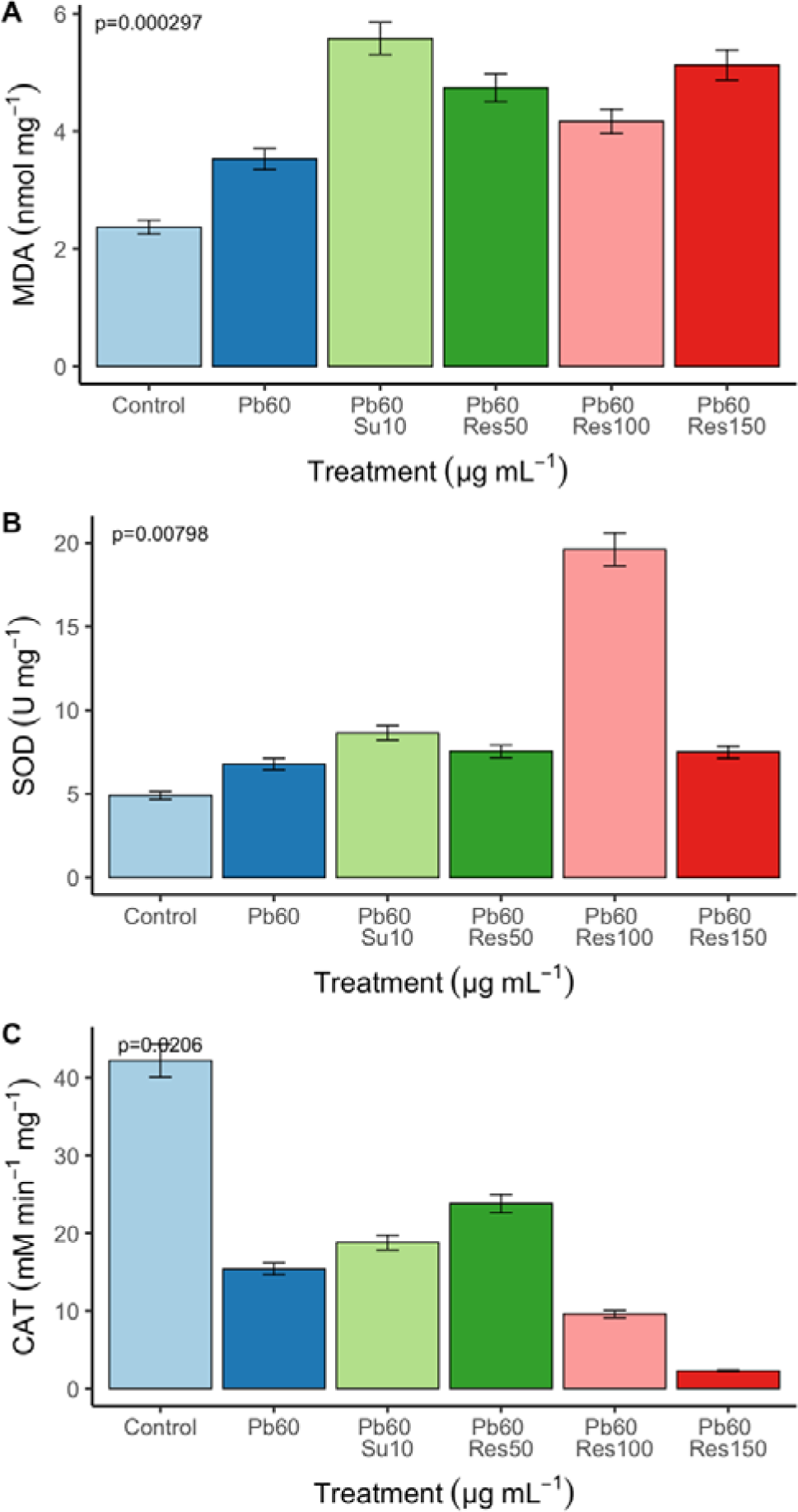
Oxidant/antioxidant status in *D. melanogaster* following 28 days of exposure to Pb, resveratrol, and succimer. Data are expressed as Mean ± SEM, with significance denoted as p < 0.05. Pb: lead concentration (mg/L), Su: succimer dosage (mg/kg), Res: resveratrol dosage (mg/kg).

## Discussion

Lead (Pb) toxicity predominantly exerts its effects at the molecular and cellular levels by inducing oxidative damage, particularly through processes like lipid peroxidation. This leads to a compromise in the body’s antioxidant defense systems, elevating levels of reactive oxygen species (Romero *et al*., 2014; Moneim, 2016; Mohamed *et al*., 2016; Balali-Mood *et al*., 2021). In the current investigation, exposure of adult Harwich strain *Drosophila melanogaster* to 60 mg/L Pb resulted in significant reductions in various biological indicators, including egg-laying capacity, eclosion rate, filial generation output, climbing ability, and lifespan. Notably, the adverse effects on these indicators were markedly alleviated by different concentrations of resveratrol.

Previous studies have reported the negative impact of lead acetate exposure on reproduction in *Drosophila*, aligning with our findings (Stamenkovic-Radak *et al.,* 2007; Mathew and Krishnamurthy, 2018). A common consequence of Pb exposure is the reduced number of eggs laid by female flies, possibly attributed to genotoxic effects leading to mutations or chromosomal abnormalities affecting egg viability and development. Additionally, interference with calcium homeostasis, as lead mimics calcium and disrupts its balance, can result in aberrations in egg maturation and release. The reduced hatching ability of pupae in the Pb-treated group may stem from toxic effects on developing embryos, as well as interference with metabolic processes and disruption of oxidative balance. Similar observations were reported by Lopez *et al*. (2015) for *D. melanogaster* exposed to green tea polyphenols and by Peterson *et al*. (2019) in response to lead acetate exposure. A significant reduction in filial generation output across five generations in flies exposed to lead suggests a potential link to mitochondrial dysfunction, impacting cell health, particularly those involved in reproduction. This reduction can significantly influence population dynamics, sustainability, and genetic diversity, affecting the population’s adaptability to changing environmental conditions. This parallels findings from studies on *Drosophila*’s reproductive fitness involving other heavy metals like Cadium (Hu et al., 2019; Yung et al., 2020; Kumari and Hena, 2022).

Exposure to Pb led to a notable decrease in the reproductive capacity of fruit flies, potentially stemming from oxidative stress, genotoxicity, and apoptosis. However, the addition of resveratrol to the diet of lead-exposed *Drosophila* significantly enhanced reproductive performance. This improvement is likely attributed to resveratrol’s ability to scavenge free radicals and prevent lipid peroxidation, thereby protecting reproductive tissues. These results are consistent with studies on other organisms, such as *Nothobranchius guentheri* (Lee et al., 2018) and recent findings for *D. melanogaster* (Igharo et al., 2023). The climbing ability, frequently utilized as an indicator of neuromuscular function in *Drosophila*, was diminished in flies exposed to lead. This reduction implies an inhibition of neuromuscular function resulting from cellular damage. Notably, male flies exhibited higher climbing activities compared to females. Hirsch *et al*. (2012) demonstrated that chronic exposure of *Drosophila* to lead decreased the neuromuscular activity of the fruit fly. Concurrently, a decrease in the lifespan of flies was observed upon lead exposure, potentially stemming from oxidative stress, DNA damage, and apoptosis. This aligns with the findings of Yung *et al*. (2020), who reported a decline in the lifespan of *D. melanogaster* exposed to Cd. The supplementation of 100 mg/kg and150 mg/kg resveratrol significantly enhanced the climbing abilities and lifespan of *D. melanogaster* exposed to lead. This improvement is likely attributed to resveratrol’s capacity to protect against oxidative stress by upregulating the phosphatase and tensin homolog (PTEN). PTEN upregulation decreases Akt phosphorylation, leading to the upregulation of antioxidant enzymes and a reduction in apoptosis (Gao *et al.,* 2016; Koushki *et al.,*2017; Abolaji *et al.,* 2018; Meng *et al.,* 2020). This is similar to the finding of Wu *et al*. (2018), who reported grape skin extract and resveratrol improved the lifespan of PD flies.

Uric acid (Ur) serves as a metabolic byproduct of purine excretion in insects, including *D. melanogaster*, functioning as an antioxidant capable of scavenging reactive oxygen species (ROS) and shielding cells from oxidative damage (Sautin and Johnson, 2008). In this study, the Ur level significantly increased in all treatment groups compared to the control, suggesting a potential protective mechanism against oxidative stress. Nevertheless, elevated uric acid levels can have adverse effects on the body, such as the formation of urate crystals, leading to metabolic disorders (Cleophas et al., 2013; Kimura et al., 2021). Reduced expression of the Urate oxidase gene in *Drosophila* resulted in elevated UA levels, aligning with the findings of Saini et al. (2020), who observed an increase in Uric acid in *D. melanogaster* larvae exposed to Cadmium and Mercury. Interestingly, in the Pb 60 mg/kg + 150 mg/kg Res group, the level of Ur was significantly increased. This could be attributed to resveratrol’s potential to enhance the activity of xanthine oxidase, an enzyme involved in uric acid synthesis (Agbadua et al., 2022). In contrast, Zhang et al. (2021) reported that resveratrol significantly reduced uric acid blood levels in C57BL/6J mice, indicating potential variations in responses across different organisms.

Creatinine, a metabolic waste product of creatine produced by muscles and eliminated through the kidneys, serves as an indicator of kidney function (Bhavsar and Kottgen, 2013). Surprisingly, the creatinine level was significantly reduced in lead-exposed flies, suggesting potential impairment in Malpighian tubule function. This contradicts the findings of Arslan et al. (2022), who reported an increase in creatinine in Japanese quail exposed to lead. The observed differences may be attributed to variations in organism classes. Resveratrol at 50 mg/kg significantly increased the level of creatine in lead-exposed flies, implying its potential protective effects against oxidative damage caused by lead toxicity. This aligns with the study by Xiao et al. (2016), where resveratrol prevented lead-induced kidney damage in mice, as evidenced by decreased levels of creatinine and other kidney function markers.

The levels of TB and CB in treatment groups were significantly higher when compared the control. Increased bilirubin production in flies co-exposed to Pb might have resulted from defects in hepatic uptake or conjugation, which may be due to destructions caused by Pb. High levels of TB and CB are generally associated with liver dysfunction and oxidative stress in various organisms. Lead acetate exposure has been shown to induce oxidative stress in *D. melanogaster* resulting in increased production of ROS and damage to cellular biomolecules. This oxidative stress can lead to liver damage and dysfunction, which may contribute to elevated levels of TB and CB (Kundrapu and Noguez, 2017) Dobrakowski, et al. (2016) also reported an increased level in TB and CB in male workers exposed to lead. The co-treatment of resveratrol at a concentration of 150 mg/kg significantly reduced the total and conjugated bilirubin, suggesting a protective effect on the fat bodies of the fruit flies.

The level of AST was significantly increased in flies exposed to lead (p < 0.05). Aspartate aminotransferase is an enzyme found mainly in the liver, heart, and skeletal muscles, and is involved in amino acid metabolism. Similar to the finding of this study, Mazumdar and Goswami (2013) reported a significant increase in the levels of AST in plastic industry workers chronically exposed to lead in India. The increased level of AST may indicate hepatocellular injury leading to enzyme leakage into the bloodstream. The addition of 10 mg/kg Su and 100 mg/kg res significantly reduced the level of AST, suggesting a protective role. Hammoud and Shalaby (2019), investigated the protective effects of resveratrol on Aluminum-induced toxicity in rats. The result showed that treatment with resveratrol significantly reduced the levels of AST in lead-exposed rats. Alkaline phosphatase (ALP) is an important enzyme in various metabolic processes, including liver function, and DNA synthesis. The enzyme is also considered as a marker for liver function in insects. When exposed to lead acetate, *Drosophila melanogaster* showed significant decrease in the level of ALP. This decrease in ALP activity may be attributed to lead-induced oxidative stress, which can lead to lipid peroxidation and damage to cell membranes. Several studies have demonstrated the inhibitory effect of heavy metals on ALP activity in different organisms, including *D. melanogaster* (Treviño *et al*. (2015); Hammoud and Shalaby, (2019). Resveratrol at 150 mg/kg significantly mitigated the effect of lead by increasing the level of ALP. This suggests that resveratrol may improve fat body function, which is corroborated by several studies (Schmatz *et. al..,* 2012; Faghihzadeh *et al*., 2015). A significant increase in co-enzyme Zn was observed in the Pb treated group, with the co-treatment group of 10 mg/kg Su having the highest level of Zn when compared to the control. Zinc has been reported to be protective against lead toxicity in various organisms, including insects (Goyer, 1997; Kazi *et al*., 2008; Lyu *et al*. 2019; Wang *et al*., 2019; Wani *et al.,* 2021).

The electrolytes, Na (p > 0.05) and Cl (p < 0.05) were increased in the Pb treated group when compared with the control. Sodium is an essential electrolyte critical in maintaining proper hydration, acid-base balance, and nerve and muscle function in animals, including insects. However, an excessive sodium level in insects can cause osmoregulatory stress, resulting in increased water loss, altered feeding behavior, and reduced lifespan (Lihoreau *et al*., 2012). Chlorine, on the other hand, is essential for many physiological functions in insects, such as, osmoregulation (Dow, 2017). The addition of resveratrol at 100 mg/kg and 150 mg/kg significantly reduced the level of Na and Cl respectively, which could imply an increase in the excretion rate of the electrolytes. Several studies in rats/mice reported an increase in Na excretion in Resveratrol treated groups (Silan, 2008; Atmaca *et al.,* 2014; Jia, *et al.,* 2020). Lead is known to interfere with the normal functioning of calcium ions in cells and disrupt calcium homeostasis (Ferreira de Mattos, *et al.,* 2017). This can lead to various consequences, including oxidative stress and apoptosis (programmed cell death). Calcium signaling is crucial for various cellular processes, including those involved in egg development and maturation. There was reduction (p > 0.05) in the level of Ca in lead-exposed flies when compared to the control group. This could imply a blockage in the Ca^2+^ channel in the cells responsible for releasing calcium in the lymph of the fly. Pb2+ exposure produced changes in the regulation of [Ca2+]i during impulse activity in *Drosophila* larva via Pb2+-dependent reduction in PMCA activity (He, *et al.,* 2009). A significant decrease was observed in the potassium (K) level in the group exposed to Pb only. This effect was significantly rescued in the 60 mg/L Pb + 150 mg/kg res group, indicating the ameliorative role of resveratrol. Potassium is an important electrolyte in insects, involved in various physiological processes including muscle contraction, nerve function, and osmoregulation (Dow, 2017). This could be the reason for the significantly low climbing abilities of the fruit fly observed in this study.

Exposure to heavy metals have been shown to induce oxidative stress in *D. melanogaster,* resulting in increased levels of malondialdehyde (MDA) as a marker of lipid peroxidation (Liu *et al.,* 2020; Yang *et al*., 2022). The level of MDA was significantly higher when compared to the control but significantly lower when compared with the resveratrol treated groups (p < 0.05). Elevated MDA levels suggest that the cell membrane has been damaged by Pb and lipid peroxidation has occurred, which can lead to cell dysfunction and death. Like the finding of this study, Aslan *et al*. (2022), also reported an increase in MDA level in quail birds exposed to Pb. The addition of resveratrol did not mitigate oxidative stress-induced cellular damage. It could be that the concentration of resveratrol used in this study was insufficient to produce an antioxidant effect on oxidative stress caused by the increase in MDA level or the exposure time was inadequate. This contradicts studies suggesting that resveratrol has potent antioxidant properties and can protect against oxidative stress-induced damage caused by an increased MDA (Farkhondeh, *et al.,* 2020; Rao, *et al.,* 2021; Hu *et al.,* 2022).

The level of SOD significantly increased when exposed to lead alone compared to the control group. Superoxide dismutase is a key antioxidant enzyme that plays a crucial role in protecting cells from oxidative damage by catalyzing the conversion of superoxide radicals to hydrogen peroxide and oxygen (Kobayashi, *et al.,* 2019). The increase observed in this study indicates that the fly is mounting an antioxidant response to combat the oxidative stress induced by lead exposure. A significant increase was observed in groups co-treated with resveratrol, with 60 mg/L Pb + 100 mg/kg Res recording the highest level of SOD activity. Increase in SOD activity due to overexpression has been shown to have increase the lifespan of *Drosophila*, which could possibly be the reason for the increased lifespan observed in this study when co-treated with resveratrol. This is similar to the reports of Shen, *et al*. (2013), Hoffman, *et al*. (2023), Nurdyansyah, *et al*. (2023). CAT activity was significantly reduced in all treated group when compared to the control (p < 0.05), with 60 mg/L Pb + 150 mg/kg Res having the least level of CAT. Catalase is crucial in detoxifying hydrogen peroxide and other harmful reactive oxygen species (ROS) in cells (Nandi *et al*. 2019). The decrease observed may be due to the generation of ROS, indicating an impaired antioxidant defense mechanism. The concentrations of resveratrol however, did not ameliorate the effect of lead on the CAT activity of the fruit flies, which might suggest an inhibitory effect of ions such as Zinc (Atli, *et al.,* 2006). It could also suggest that the doses of resveratrol used was either too high or too low to have elicit a positive effect on the CAT activity, or it could be that the time of exposure was not sufficient enough, or the increase in SOD level observed in this study, which leads to an increase in the dismutation of O2⎯ and H2O2 could have led to the decrease of CAT activity as reported by Sullivan-Gunn and Lewandowski (2013) and Virgolini and Aschner (2021). The stimulation of SOD activity and reduction in CAT activity by Pb observed in this study is in agreement with the reports of Wang *et al*. (2015) for Cd.

## Conclusion

Lead toxicity in *D. melanogaster* is attributed to an imbalance between reactive oxygen species (ROS) and the antioxidant defense system, decreasing various biological and biochemical parameters. Resveratrol has shown to have a potential therapeutic effect against lead-induced toxicity in *Drosophila* melanogaster, probably by its antioxidant and anti-inflammatory properties. These results suggest the possible use of resveratrol as an affordable and safe therapeutic agent for preventing lead toxicity in humans. The safety of resveratrol in flies indicates its potential for treating neurodegenerative, hepatic, and renal diseases. However, further studies are required to fully comprehend the mechanisms underlying the protective effects of resveratrol against lead toxicity and to evaluate its safety and efficacy as a therapeutic agent for lead toxicity.

## Author contributions

Conceptualization: S.M.H and R.A.; data curation: U.F.O., M.T.J., H.M., S.M.H. and R.A.; formal analysis and investigation: U.F.O., M.T.J., H.M., S.M.H. I.S.N. and R.A. writing— original draft, R.A; writing, reviewing and editing: R.A., D.M.S. and I.S.N.; supervision: I.S.N., S.M.H and R.A. Fund awardees: R.A., D.M.S. and I.S.N.

## Acknowledgements

The authors would like to express their sincere gratitude to Mal. S. Muhammad for his valuable assistance during the lab work, and to Dr. M.A. Chia for his invaluable contribution in proofreading this article. His expertise and attention to detail greatly improved the quality of the manuscript.

